# 3D-MASNet: 3D Mixed-scale Asymmetric Convolutional Segmentation Network for 6-month-old Infant Brain MR Images

**DOI:** 10.1101/2021.05.23.445294

**Authors:** Zilong Zeng, Tengda Zhao, Lianglong Sun, Yihe Zhang, Mingrui Xia, Xuhong Liao, Jiaying Zhang, Dinggang Shen, Li Wang, Yong He

**Author notes:** Correspondence (Y.H.) or (T.Z).

## Abstract

Precise segmentation of infant brain MR images into gray matter (GM), white matter (WM), and cerebrospinal fluid (CSF) are essential for studying neuroanatomical hallmarks of early brain development. However, for 6-month-old infants, the extremely low-intensity contrast caused by inherent myelination hinders accurate tissue segmentation. Existing convolutional neural networks (CNNs) based segmentation models for this task generally employ single-scale symmetric convolutions, which are inefficient for encoding the isointense tissue boundaries in baby brain images. Here, we propose a 3D mixed-scale asymmetric convolutional segmentation network (3D-MASNet) framework for brain MR images of 6-month-old infants. We replaced the traditional convolutional layer of an existing to-be-trained network with a 3D mixed-scale convolution block consisting of asymmetric kernels (MixACB) during the training phase and then equivalently converted it into the original network. Five canonical CNN segmentation models were evaluated using both T1- and T2-weighted images of 23 6-month-old infants from iSeg-2019 datasets, which contained manual labels as ground truth. MixACB significantly enhanced the average accuracy of all five models and obtained the most considerable improvement in the fully convolutional network model (CC-3D-FCN) and the highest performance in the Dense U-Net model. This approach further obtained Dice coefficient accuracies of 0.931, 0.912, and 0.961 in GM, WM, and CSF, respectively, ranking first among 30 teams on the validation dataset of the iSeg-2019 Grand Challenge. Thus, the proposed 3D-MASNet can improve the accuracy of existing CNNs-based segmentation models as a plug-and-play solution that offers a promising technique for future infant brain MRI studies.

## 1. Introduction

The accurate tissue segmentation of infant brain magnetic resonance (MR) images into gray matter (GM), white matter (WM), and cerebrospinal fluid (CSF) are essential for researchers to chart the normal and abnormal early brain development of cortical regions, white matter connections, and wiring topologies (Cao et al., 2017; Hazlett et al., 2017; Wang et al., 2019a; Wen et al., 2019; Xu et al., 2019; Zhao et al., 2019). Notably, the tissue segmentation of 6-month-old infants is the biggest challenge in baby brain segmentation tasks due to the isointense phase in which the intensity distributions of GM and WM voxels become dramatically overlapped in the cortical regions (Fig. 1). The effective manual annotation, which is guided by longitudinal tracking of brain images with high tissue contrast in the latter children period (Wang et al., 2019b), is limited by the extremely high labor costs, the requirement of specialized expert knowledge (almost one week per image for an experienced neuroradiologist) and high inter- and intra-rater variations (Makropoulos et al., 2018). Developing fast, automatic, and accurate brain segmentation approaches is a crucial and ongoing goal for MR images of infants at 6 months of age (Sun et al., 2021; Wang et al., 2019b).

**Figure 1.**
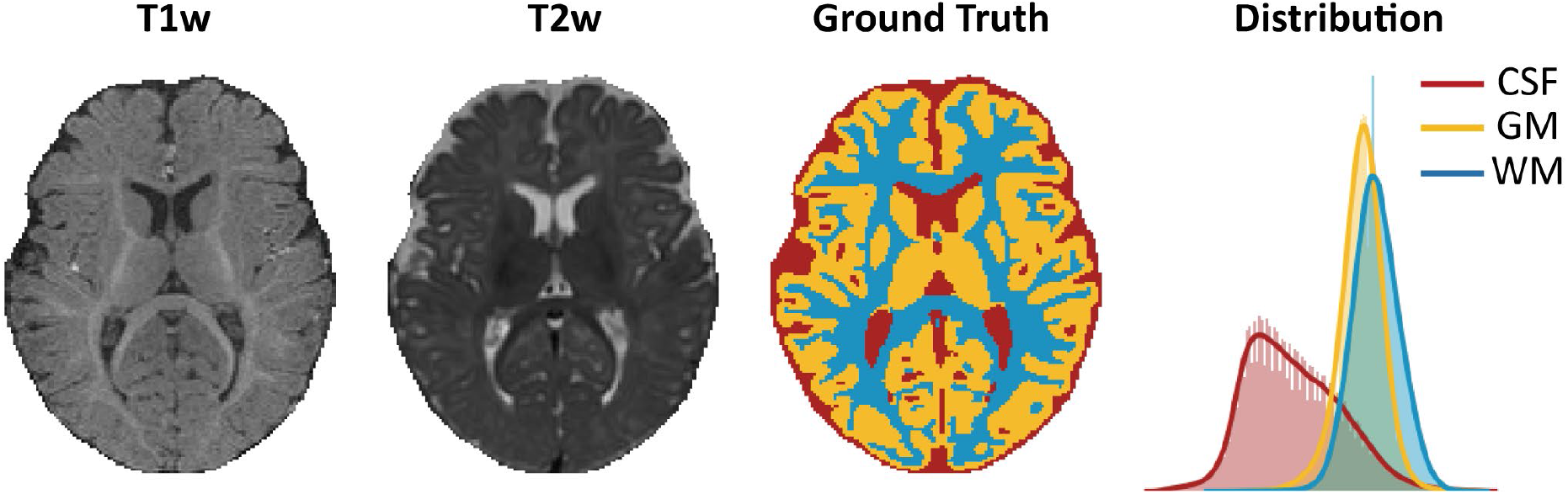
Data of a 6-month-old infant from the training set in iSeg-2019. The isointense brain appearance of an axial slice in T1-weighted (T1w) and T2-weighted (T2w) images. An axial view of the manual segmentation label (ground truth) and the corresponding brain tissue intensity distribution of the T1w image (distribution).

### 1.1. Convolutional neural networks (CNNs) based methods become the mainstream

In the past years, many efforts have been made for the segmentation task of 6-month-old infant brain MR images. Generally, emerging CNNs-based segmentation methods that exhibiting faster computational speed and higher accuracy than conventional atlas-based (Wang et al., 2014; Wang et al., 2012) or machine learning based methods (Sanroma et al., 2018; Wang et al., 2015; Wang et al., 2018a; Wang et al., 2014) become the mainstream. A typical example is that seven of the eight top teams in iSeg-2017 challenge has utilized CNNs to segment infant brain tissues.

Current CNNs-based approaches for infant brain segmentation are usually variants of canonical FCN (Long et al., 2015) and U-Net (Ronneberger et al., 2015) architecture. By adjusting or adding specific connectional pathways within or across neural layers on classical CNNs models, these approaches enhance the extraction and fusion of the semantic information in multimodal features to counteract the noisy and isointense tissues boundaries in 6-month-old infant brain images (Bui et al., 2019; Dolz et al., 2020; Dolz et al., 2019; Nie et al., 2019; Nie et al., 2016; Wang et al., 2018b; Wang et al., 2020; Zeng and Zheng, 2018; Zhang et al., 2015). Specifically, Bui et al. improved densely connected network (DenseNet) (Huang et al., 2017) by concatenating fine and coarse feature maps from multiple densely connected blocks and won the iSeg-2017 competition (Bui et al., 2019). Dolz et al. proposed a semi-dense network by directly connecting all of the convolutional layers to the end of the network (Dolz et al., 2020) and further extended it into a HyperDenseNet by adding dense connections between multimodal network paths (Dolz et al., 2019). Similarly, Zeng et al. modified the classical U-Net network by constructing multi-encoder paths for each modality to effectively extract targeted high-level information (Zeng and Zheng, 2018). Wang et al. designed a global aggregation block in the U-Net model to consider global information in the decoder path of feature maps (Wang et al., 2020). Interestingly, inspired by the superiority of DenseNet and U-Net, the densely connected U-Net (DU-Net) model with a combination of these two types of networks was proposed for both tissue segmentation and autism diagnosis (Wang et al., 2018b).

### 1.2. Improvements from fine-grained convolution kernel designs are underestimated

Although great efforts have been made, the above CNN-based segmentation models have several limitations. First, the image appearance of 6-month-old infant brain MR images is quite noisy (Li et al., 2019; Mostapha and Styner, 2019) which makes the effective feature extraction difficult for the traditional convolution kernel design in previous works. Adopting enhanced convolution kernel designs (Ding et al., 2019; Li et al., 2020; Zhang et al., 2022) that emphasizes key features in the skeleton center of kernels may facilitate feature extractions throughout the network. Second, the voxel-wise fuzzy tissue boundaries in infant brain images are constrained by the anatomical morphology of gyrus at large spatial scales (Wang et al., 2018a). Although previous infant segmentation approaches try to fuse multi-scale features by skip-connections in variants of FCN and U-Net, they overlook capturing rich multi-scale features in kernel space, which contains more stable and homogeneous semantic information than features between layers (Fan et al., 2019). Third, all these studies focused on modifications of network layouts which need seasoned expertise experience, time-consuming hyperparameter tuning, and may also bring excessive graphics processing unit (GPU) burdens (Dolz et al., 2019; Wang et al., 2020) and architecture incompatibility. Recent CNN studies move eyes on building architecture-independent designs such as SE blocks (Hu et al., 2018), or automatically configuring models such as nnU-net (Isensee et al., 2021), which requires neither rare expert knowledge nor expensive manual interventions.

### 1.3. Our contribution

Our goal is to obtain a CNN-based building block for 6-month-old infant brain image segmentation which is 1) with fine-grained kernel designs to enhance the representation and abundance of features; 2) transplantable in up-to-date segmentation models in a plug-and-play way; 3) without much additional hyperparameter tuning or computational burden. To this end, we construct a 3D mixed-scale asymmetric segmentation network (3D-MASNet) framework by embedding a well-designed 3D mixed-scale asymmetric convolution block (MixACB) into existing segmentation CNNs for 6-month-old infant brain MR images (Fig. 2). The MixACB design is comprised by 1) four parallel 3D convolutional layers including a symmetric kernel (*d* × *d* × *d*) and three asymmetric 2D kernels (1× *d* × *d*, *d* × 1 × *d*, *d* × *d* ×1) (Fig. 3A), respectively; 2) multiple groups on input feature maps with different kernel sizes (Fig. 3B) independently; 3) parameter fusion for each MixACB after the training process to lower inference-time computations compare to the original network. We first evaluated the effectiveness of the MixACB on five canonical CNN networks using the iSeg-2019 training dataset. We next compared the performance of our method with that of top-4 approaches proposed in the MICCAI iSeg-2019 Grand Challenge on the iSeg-2019 validation dataset. The experimental results revealed that the MixACB significantly improved the segmentation accuracy of various CNNs, among which DU-Net (Wang et al., 2018b) with MixACB achieved the best-enhanced average performance and obtained the highest Dice coefficients of 0.931 in GM, 0.912 in WM, and 0.961 in CSF, ranking first in the iSeg-2019 Grand Challenge. All codes are publicly available at https://github.com/RicardoZiTseng/3D-MASNet.

**Figure 2.**
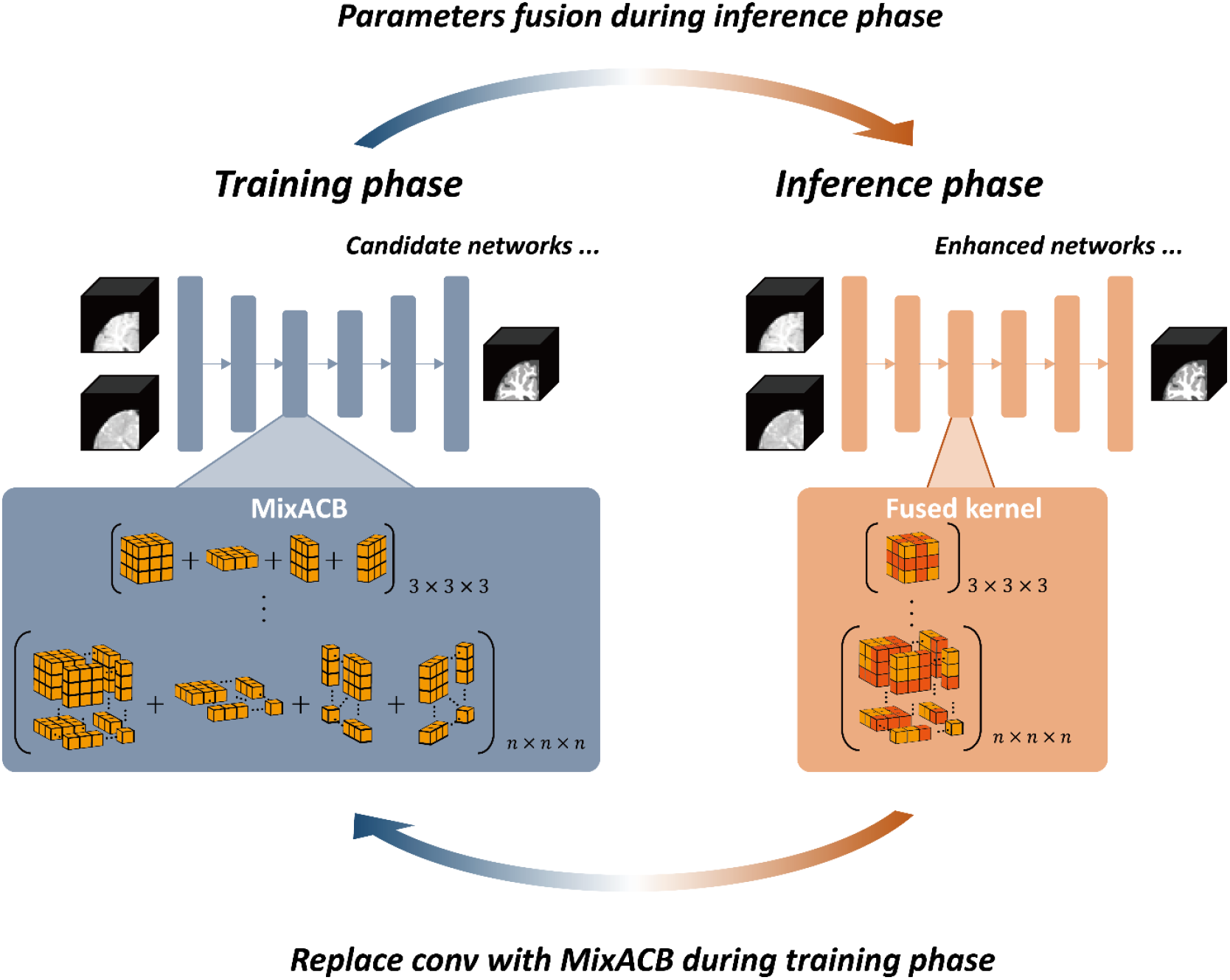
Overview of the 3D-MASNet framework. For a candidate network, we replace its traditional convolutional layers with MixACB during the training phase. Once the training process is complete, we fuse the parameters of MixACB to obtain an enhanced model containing fewer parameters after equivalent fusion.

**Figure 3.**
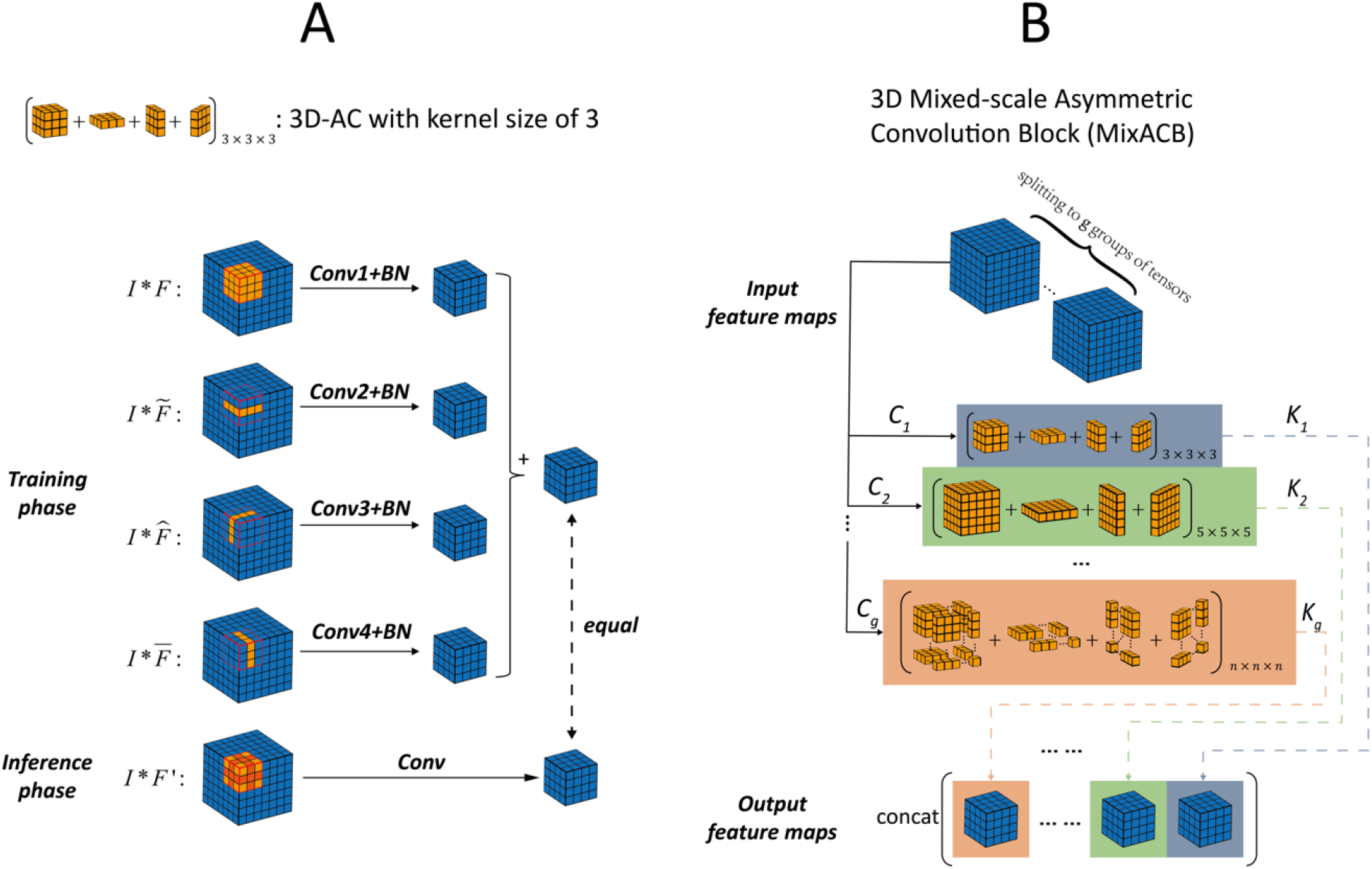
(A) Diagram of 3D-AC (taking a kernel size of 3 as an example), which has four convolutional layers during the training phase and one convolutional layer once kernel parameters have been fused during the inference phase. (B) Diagram of MixACB, which is composed of multiple 3D-ACs with different kernel sizes. MixACB splits input feature maps into several groups, applies asymmetric convolution on each group of feature maps, and then concatenates each group’s output as the output feature maps.

## 2. Methods and Implementations

### 2.1.1. Mathematical formulation of basic 3D convolution

Consider a feature map *I* ∈ ℝ^*U*×*V*×*S*×*C*^ with a spatial resolution of *U*×*V*×*S* as input and a feature map *O* ∈ ℝ^*R*×*T*×*Q*×*K*^ with a spatial resolution of *R*×*T*×*Q* as output of a convolutional layer with a kernel size of *H*×*W*×*D* and *K* filters. Then, each filter’s kernel is denoted as *F* ∈ ℝ^*H*×*W*×*D*×*C*^, and the operation of the convolutional layer with a batch normalization (BN) layer can be formulated as follows:

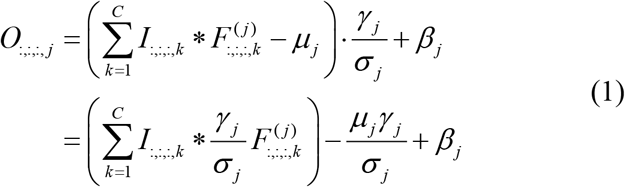

where * is the 3D convolution operator, 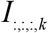 is the *k^th^* channel of the input feature map *I*, 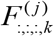 is the *k^th^* channel of the *j^th^* filter’s kernel, *μ_j_* and *σ_j_* are the channel-wise mean value and standard deviation value, respectively, *γ_j_* and *β_j_* are the scaling factor and bias term to restore the representation ability of the network, respectively.

### 2.1.2. Design of 3D asymmetric convolutions (3D-AC) during training and inference phases

3D-AC was designed behaving differently during training and inference phases (Fig. 3A). Concretely, for each kernel of each layer in the network during the training phase, a 3D-AC contains 4 parallel convolutional branches, namely one standard 3D convolution layer and three orthogonal 2D asymmetric convolutional layers (1× *d* × *d, d* ×1× *d*, *d* × *d* ×1) at kernel center for the enhancement of features along axial, sagittal and coronal directions, respectively. The input feature maps are fed into these 4 branches, and the outputs of these branches are summed to fuse the knowledge learned by these 4 independent branches. During the inference phase, 3D-AC contains one standard convolutional layer with equivalently fused kernel of the training-time 3D-AC (described in section 2.1.4), thus the input feature maps only need feed into this single branch which brining low inference computations.

### 2.1.3. Constructing MixACB by multiple 3D-ACs with varying kernel scales

To process the input feature map at different scales of detail, we propose the MixACB by mixing multiple 3D-ACs with different kernel sizes, as illustrated in Fig. 3B. Notably, since we used the 3D-AC to strength the core skeleton part of the convolutional kernel, thus the kernel size of 3D-AC must be odd, such as 3, 5, 7. Since directly adopting multiple 3D-ACs on all feature maps then concatenating outputs will dramatically increase the models’ parameters and computations, we leverage the grouped convolution approach by splitting original input feature maps into groups and apply 3D-AC independently in each input feature map’s group. Assume that we split the input feature maps into *g* groups of tensors such that their total number of channels is equal to the original feature maps’ channels: *C*_1_ + *C*_2_ +⋯ + *C_g_* = *C* with *C*_1_ > *C*_2_ >⋯ > *C_g_*; similarly, the output feature maps also have *g* groups: *K*_1_ + *K*_2_ +⋯ + *K_g_* = *K* with *K*_1_ >*K*_2_ >⋯ > *K_g_*. We denote *I*^<*i*>^ ∈ ℝ^*U*×*S*×*C_i_*^ as the *i^th^* group of input, 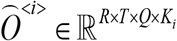 as the MixACB’s *i^th^* group output, and *F*’^<*i*>^ ∈ ℝ^*H_i_×W_i_×D_i_×C_i_*^ as the equivalent kernel of the *i^th^* group of the 3D-AC whose equivalent kernel size is *H_i_*×*W_i_*×*D_i_*. Thus, we have following equations:

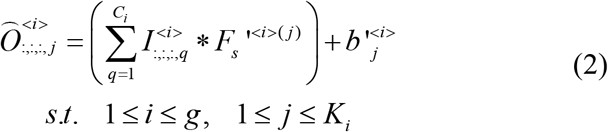

The final output of MixACB is the concatenation of all groups’ outputs:

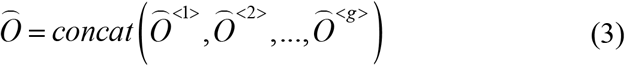

In this paper, we only split the input and output feature maps into 2 groups, and define the mix ratio as the ratio between *C*_1_ and *C*_2_. For simplicity, the ratio between *K*_1_ and *K*_2_ is set to be equal to the mix ratio. We set a kernel size of 3 for the 1^st^ group of 3D-AC and a kernel size of 5 for the 2^nd^ group of 3D-AC, with the mix ratio set to 3:1. In this situation, the number of FLOPs (floating point operations) required for inference-time MixACB is 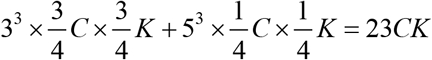, which is smaller than that (3^3^ × *C* × *K* = 27*CK*) for standard convolutions with kernel size of 3.

### 2.1.4. Equivalently fusing kernel of each 3D-AC inside MixACB

Once the training process of 3D-MASNet is completed, we equivalently fused the kernels of each 3D-AC inside the MixACB to retain the same output results as the original network. Due to the additivity of convolutional kernels, the kernels of 3D-AC’s four branches can be fused to obtain an equivalent kernel in a 3D convolutional layer to produce the same output, which can be formulated as the following equation:

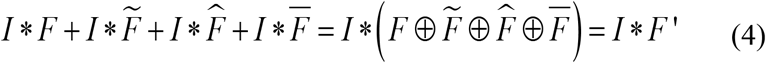

where *I* is an input feature map, 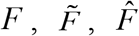 and 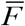 are the 4 branches’ kernels of 3D-AC. ⊕ is an elementwise operator that performs parameter addition on the corresponding positions, and *F*′ is the equivalent fused kernel of the 4 branches’ kernels.

Here, we took a kernel size of 3 as an example. We first fused the BN parameters into the convolutional kernel term and bias term following Eq. (1). Then, we further fused the four parallel kernels by adding the asymmetric kernels onto the skeletons of the cubic kernel. Formally, we denote *F*’^(*j*)^ as the *j^th^* filter at the 1 × 3 × 3, 3 × 1 × 3 and 3 × 3 × 1 layer, respectively. Hence, we obtain the following formulas:

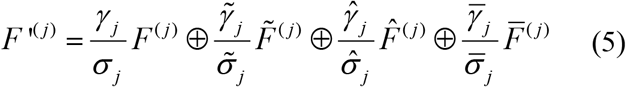

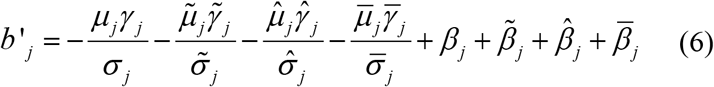

Then, we can write any output of *j^th^* filter as:

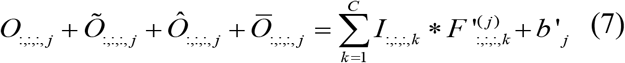

where 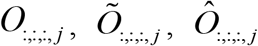 and 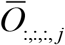 are the outputs of the original 3 × 3 × 3, 1 × 3 × 3, 3 × 1 × 3 and 3 × 3 × 1 branch, respectively.

### 2.2. Candidate CNNs for the evaluation of the MixACB on 6-month-old infant brain image segmentation

We choose five representative networks to evaluate the effectiveness of the 3D-MixACB in improving the segmentation performance, including BuiNet (Bui et al., 2019), 3D-UNet (Çiçek et al., 2016), convolution and concatenate 3D fully convolutional network (CC-3D-FCN) (Nie et al., 2019), non-local U-Net (NLU-Net) (Wang et al., 2020) and DU-Net (Wang et al., 2018b). Notably, these five networks are either variants of the U-type architecture (3D U-Net, NLU-Net, and DU-Net) or the FCN-type architecture (BuiNet and CC-3D-FCN) and encompass major CNN frameworks in infant brain segmentation. After replacing their original convolution layers with the 3D-MixACB design, we followed the training configurations set in the candidate CNN’s release codes (Table 1) and adopted the Adam optimizer to update these models’ parameters. Except for the CC-3D-FCN, which used the Xavier algorithm (Glorot and Bengio, 2010) to initialize network weights, all other networks adopted the He initialization method (He et al., 2015). The configuration parameters are as follows:

1. BuiNet adopted four dense blocks consisting of four 3 × 3 × 3 convolutional layers for feature extraction. Transition blocks were applied between every two dense blocks to reduce the feature map resolutions. 3D up-sampling operations were used after each dense block for feature map recovery, and these upsampled features were concatenated together. (2) 3D-UNet has 4 levels of resolution, and each level adopts one 3 × 3 × 3 convolution, which is followed by BN and a rectified linear unit (ReLU). The 2 × 2 × 2 max pooling and the 2 × 2 × 2 transposed convolution, each with a stride of 2, are employed for resolution reduction and recovery. Feature maps of the same level of both paths were summed. (3) CC-3D-FCN used 6 groups of 3 × 3 × 3 convolutional layers for feature extraction, in which the 2 × 2 × 2 max pooling with a stride of 2 was adopted between two groups of layers. The 1 × 1 × 1 convolution with a stride of 1 was added between two groups with the same resolution for feature fusion. (4) DU-Net used 7 dense blocks to construct the encoder-decoder structure with 4 levels of resolution and leveraged transition down blocks and transition up blocks for down-sampling and up-sampling, respectively. Unlike the implementations in (Wang et al., 2018b), the bottleneck layer is introduced into the dense block to constrain the rapidly increasing number of feature maps, and the transition down block consisted of two 3 × 3 × 3 convolutions, each followed by BN and ReLU. In addition, we used the 1 × 1 × 1 convolution followed by a softmax activation function in the last layer. (5) NLU-Net leveraged five different kinds of residual blocks to form the U-type architecture with 3 levels of resolution. BN with the ReLU6 activation function was adopted before each 3 × 3 × 3 convolution. The global aggregation block replaced the two convolutional layers of the input residual block to form the bottom residual block for the integration of global information.

**Table 1.** Training strategy of each candidate network

| Candidate Network | Training Batch Size | Training/Inference Patch Size | Learning Rate Schedule |
| --- | --- | --- | --- |
| BuiNet | 4 | 64 | Train for 20,000 iterations. The initial learning rate is set to $2e-4$ and is decreased by a factor of 0.1 every 5000 iterations. |
| 3D-UNet | 10 | 32 | Train for 80 epochs for a total of 5000 patches that are randomly extracted per epoch. The learning rate is decreased every 20 epochs and is set to $3e-4$, $1e-4$, $1e-5$ and $1e-6$. Train for 80 epochs for a total of 5000 patches that are randomly extracted per epoch. The learning rate is decreased every 20 epochs and is set to $3e-4$, $1e-4$, $1e-5$ and $1e-6$. |
| CC-3D-FCN | 10 | 32 | The same as 3D-UNet. |
| NLU-Net | 5 | 32 | Train for 80 epochs for a total of 5000 patches that are randomly extracted per epoch. The learning rate is set to $1e-3$. |
| DU-Net | 16 | 32 | The cosine annealing strategy with a maximum learning rate of $3e-4$ and a minimum learning rate of $1e-6$ is adopted. The model is trained for 500 epochs and a total of 1000 patches are randomly extracted at each epoch. |

We fed the same multimodal images into these five networks and employed the same inference strategy. We extracted overlapping patches of the same size as that used during the training phase. The overlapping step size had to be smaller than or equal to the patch length size to form the whole volume. Following the common practice in (Bui et al., 2019; Nie et al., 2019; Wang et al., 2018b; Wang et al., 2020), we set the step size to 8. Since the effect of the overlapping step size in the proposed framework remains unknown, we further evaluated it in section 3.3. Voxels inside the overlapping regions were averaged.

## 3. Experiments and Results

### 3.1. iSeg-2019 dataset and image preprocessing

Twenty-three isointense phase infant brain MRIs, including T1w and T2w images, were offered by the iSeg-2019 (http://iseg2019.web.unc.edu/) organizers from the pilot study of the Baby Connectome Project (BCP) (Howell et al., 2019). All the infants were term-born (40±1 weeks of gestational age) with an average scan age of 6.0±0.5 months. All experimental procedures were approved by the University of North Carolina at Chapel Hill and the University of Minnesota Institutional Review Boards. Detailed imaging parameters and preprocessing steps that were implemented are listed in (Sun et al., 2021). Before cropping the MR images into patches, we normalized the T1w and T2w images by subtracting the mean value and dividing by the standard deviation value.

The iSeg-2019 organizers offered the ground truth labels, which were obtained by a combination of initial automatic segmentation using the infant brain extraction and analysis toolbox (iBEAT) (Dai et al., 2013) on follow-up 24-month scans of the same baby and manual editing using ITK-SNAP (Yushkevich et al., 2006) under the guidance of an experienced neuroradiologist. The MR images of 10 infants with manual labels were provided for model training and validation. The images of 13 infants without labels were provided for model testing. The testing results were submitted to the iSeg-2019 organizers for quantitative measurements.

### 3.2. Evaluation metrics

We employed the Dice coefficient (DICE), modified Hausdorff distance (MHD) and average surface distance (ASD) to evaluate the model performance on segmenting 6-month-old infant brain MR images.

#### 3.2.1. Dice coefficient

Let *A* and *B* be the manual labels and predictive labels, respectively. The DICE can be defined as:

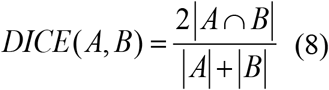

where |·| denotes the number of elements of a point set. A higher DICE indicates a larger overlap between the manual and predictive segmentation areas.

#### 3.2.2. Modified Hausdorff distance

Let *C* and *D* be the sets of voxels within the manual and predictive segmentation boundary, respectively. MHD can be defined as:

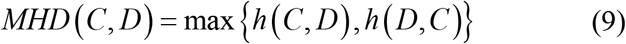

where 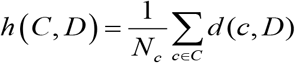, and 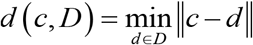 with 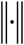 representing the Euclidean distance. We follow the calculation described in (Wang et al., 2020) by computing the average MHD based on the three different vectorization directions to obtain a direction-independent evaluation metric. A smaller MHD coefficient indicates greater similarity between manual and predictive segmentation contours.

#### 3.2.3. Average surface distance

The ASD is defined as:

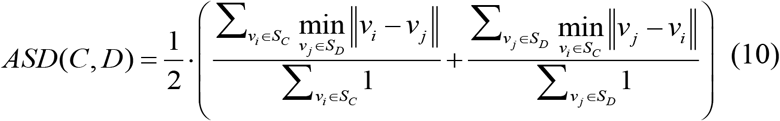

where *S_C_* and *S_D_* represent the surface meshes of *C* and *D*, respectively. A smaller ASD coefficient indicates greater similarity between cortical surfaces reconstructed from manual and predictive segmentation maps.

### 3.3. Exploring the effectiveness of the MixACB

We performed several experiments to evaluate the effectiveness of the MixACB, including 1) ablation tests on five representative segmentation networks (section 2.2); 2) comparisons with state-of-the-art approaches in iSeg-2019; 3) component analysis of MixACB; and 4) validation of the impact of the overlapping step size.

#### 3.3.1. Performance improvement on five representative CNN architectures

For a given network architecture without the MixACB design, we regarded it as the baseline model and further transformed it into a 3D-MASNet design. All pairs of the baseline models and their corresponding 3D-MASNet followed the training strategies described in Table 1. To balance the training and testing sample sizes, we adopt a 2-fold cross-validation (one fold with five random selected participants for training and the left for testing) for model evaluation on the iSeg-2019 training dataset. Table 2 and Table 3 and Fig. 4 show that the performance of all the models was significantly improved across almost all tissue types in terms of the DICE and MHD, which demonstrates the effectiveness of the MixACB on a wide range of CNN layouts. Specifically, DU-Net with the MixACB achieved the highest average DICE of 0.928 and the lowest average MHD value of 0.436; CC-3D-FCN with the MixACB gained the most considerable DICE improvement and reached a higher average DICE than that attained by BuiNet, which was a champion solution in the MICCAI iSeg-2017 grand challenge, indicating that a simple network could reach excellent performance by advanced convolution designs. Fig. 5 further provides a visual segmentation comparison between networks with and without the MixACB. The MixACB could effectively correct misclassified voxels which are indicated by red squares.

**Figure 4.**
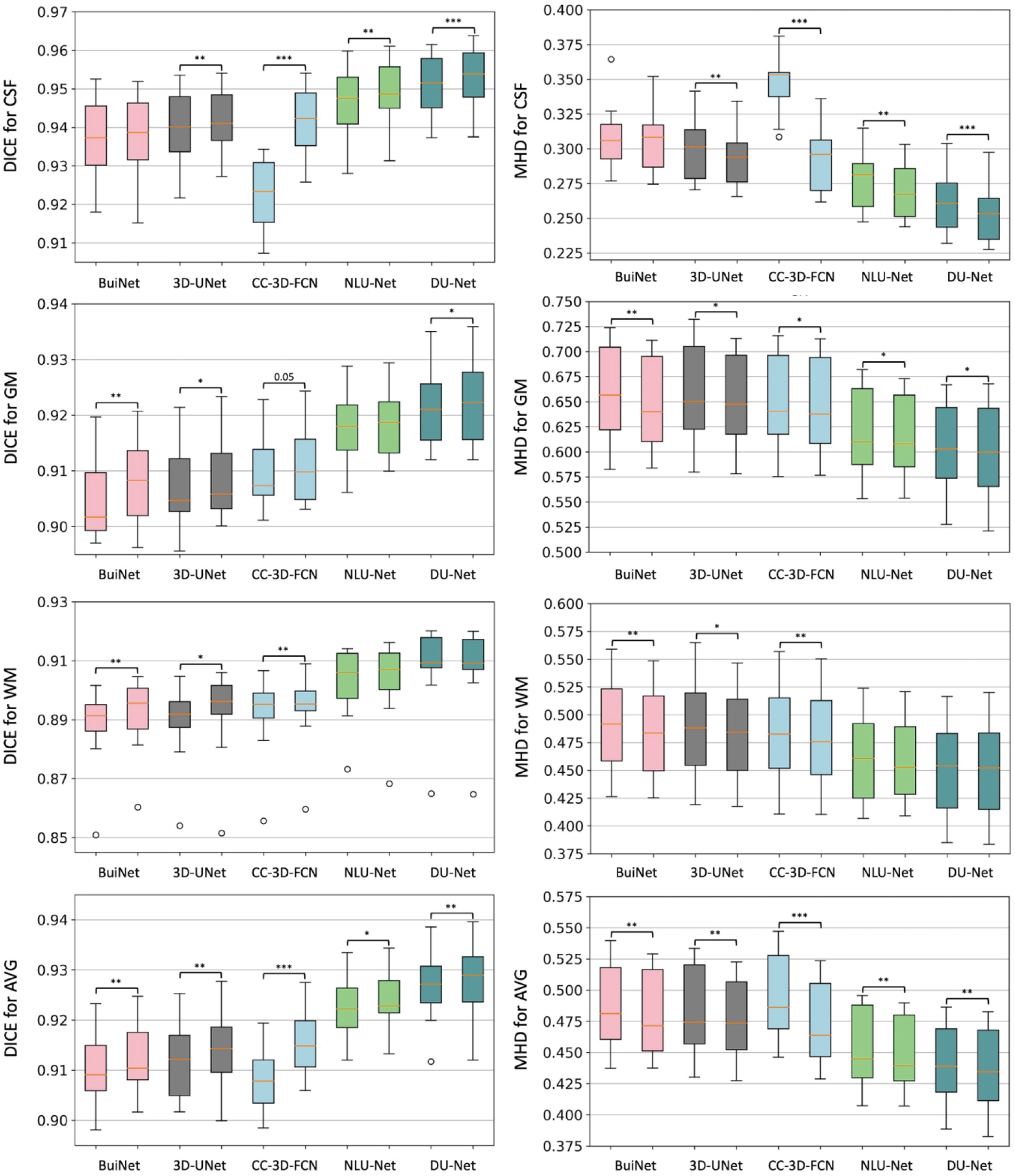
Box plot of the segmentation performance improvement on five candidate CNN architectures in the 3D-MASNet framework. The first column shows the measurement of DICE to represent the segmentation accuracy for each tissue type. The second column shows the results of MHD. In each subgraph, we use two neighbor box plots to represent a candidate model (first bar) and its corresponding 3D-MASNet (second bar). The significance of model comparison is evaluated by 2-fold cross-validation. “*” denotes that 0.01≤p<0.05, “**” denotes that 0.001≤p<0.01, and “***” denotes that p<0.001.

**Figure 5.**
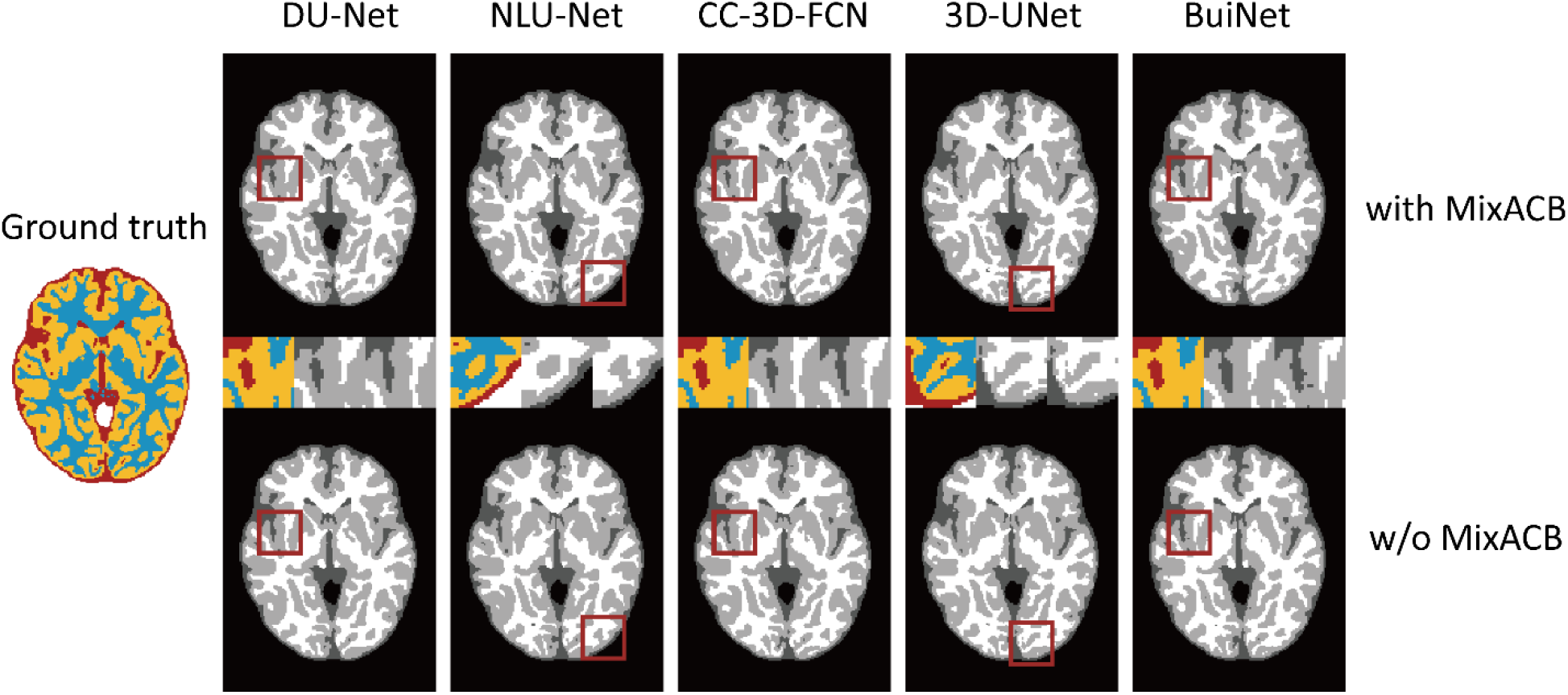
Visualization of the segmentation results on different models with (w) and without (w/o) the MixACB. The ground truth map is shown in color, and CNNs-based segmentation maps are shown in the gray scale. The regions in the red square are magnified in the middle row following an order from with MixACB to without MixACB.

**Table 2.** Ablation study performed by comparing the segmentation accuracy between different models and their corresponding 3D-MASNet in terms of DICE by 2-fold cross validation.

| Netw<br>ork | CSF |  | GM |  | WM |  | Avg |  |
| --- | --- | --- | --- | --- | --- | --- | --- | --- |
|  | Baselin<br>e | MixACB | Baselin<br>e | MixACB | Baselin<br>e | MixACB | Baselin<br>e | MixACB |
| BuiN<br>et | 0.938±0<br>.010 | 0.938±0.<br>011 | 0.905±0<br>.007 | 0.908*±<br>0.007 | 0.888±0<br>.014 | 0.892*±<br>0.013 | 0.910±0<br>.007 | 0.912*±<br>0.007 |
| 3D-UNet | 0.940±0<br>.010 | 0.942*±<br>0.008 | 0.907±0<br>.007 | 0.909*±<br>0.007 | 0.889±0<br>.014 | 0.892*±<br>0.015 | 0.912±0<br>.007 | 0.914*±<br>0.008 |
| CC-3D-FCN | 0.923±0<br>.010 | 0.942*±<br>0.008 | 0.910±0<br>.006 | 0.911±0.<br>007 | 0.892±0<br>.013 | 0.894*±<br>0.013 | 0.908±0<br>.006 | 0.915*±<br>0.006 |
| NLU<br>-Net | 0.947±0<br>.009 | 0.949*±<br>0.008 | 0.918±0<br>.007 | 0.919±0.<br>006 | 0.903±0<br>.012 | 0.904±0.<br>014 | 0.922±0<br>.006 | 0.924*±<br>0.006 |
| DU-Net | 0.951±0<br>.008 | **0.953\*±**<br>0.008 | 0.922±0<br>.007 | **0.923\*±**<br>0.007 | 0.907±0<br>.015 | **0.907±0.**<br>015 | 0.927±0<br>.007 | **0.928\*±**<br>0.008 |

Note that the best values are highlighted in bold font. “Baseline” denotes that the corresponding model adopted the standard convolutional operation; “MixACB” denotes that the corresponding model was transformed into 3D-MASNet; “\*” denotes that the difference between baseline and 3D-MASNet is statistically significant ( $p<0.05$ ).

**Table 3.** Ablation study performed by comparing the segmentation accuracy between different models and their corresponding 3D-MASNet in terms of MHD by 2-fold cross validation.

| Netw<br>ork | CSF |  | GM |  | WM |  | Avg |  |
| --- | --- | --- | --- | --- | --- | --- | --- | --- |
|  | Baselin<br>e | MixACB | Baselin<br>e | MixACB | Baselin<br>e | MixACB | Baselin<br>e | MixACB |
| BuiN<br>et | 0.308±0<br>.024 | 0.307±0.<br>023 | 0.659±0<br>.048 | 0.649*±<br>0.045 | 0.493±0<br>.043 | 0.485*±<br>0.042 | 0.487±0<br>.035 | 0.480*±<br>0.034 |
| 3D-UNet | 0.299±0<br>.022 | 0.293*±<br>0.020 | 0.658±0<br>.050 | 0.651*±<br>0.046 | 0.490±0<br>.046 | 0.485*±<br>0.042 | 0.483±0<br>.036 | 0.476*±<br>0.033 |
| CC-3D-FCN | 0.348±0<br>.022 | 0.292*±<br>0.023 | 0.649±0<br>.047 | 0.645*±<br>0.048 | 0.485±0<br>.046 | 0.480*±<br>0.046 | 0.494±0<br>.034 | 0.473*±<br>0.034 |
| NLU<br>-Net | 0.278±0<br>.022 | 0.270*±<br>0.020 | 0.619±0<br>.043 | 0.615*±<br>0.040 | 0.461±0<br>.040 | 0.460±0.<br>037 | 0.453±0<br>.032 | 0.448*±<br>0.030 |
| DU-Net | 0.261±0<br>.021 | **0.254\*±**<br>0.022 | 0.605±0<br>.046 | **0.601\*±**<br>0.047 | 0.452±0<br>.041 | **0.452±0.**<br>043 | 0.439±0<br>.032 | **0.436\*±**<br>0.034 |

Note that the best values are highlighted in bold font. “Baseline” denotes that the corresponding model adopted the standard convolutional operation; “MixACB” denotes that the corresponding model was transformed into 3D-MASNet; “\*” denotes that the difference between baseline and 3D-MASNet is statistically significant ( $p < 0.05$ ).

#### 3.3.2. Comparison with state-of-the-art methods on iSeg-2019

Since DU-Net, which was combined with MixACB, has achieved the highest accuracy among all candidate models, we compared it with methods developed by the 29 remaining teams that participated in the iSeg-2019 challenge. We employed a majority-voting strategy on 10 trained networks’ outputs to improve the model generalization.

Table 4 reports the segmentation results achieved by our proposed method and those of other teams’ methods that ranked in the top 4 on the validation dataset of the iSeg-2019. The mean DICE, MHD value and ASD value are presented for CSF, GM, and WM, representatively. Compared with other teams, our method yielded the highest DICE and lowest ASD value for the three brain tissues in the validation test of iSeg-2019, with comparable MHD values. The superior average value of the 3 types of brain tissues also indicates that our method has the best overall performance.

**Table 4.** Comparison of segmentation performance of the proposed method and the methods of the top-4 ranked teams on the 13 validation infant MRI images of iSeg-2019.

| Method (Top<br>5) | CSF |  |  | GM |  |  | WM |  |  | AVG |  |  |
| --- | --- | --- | --- | --- | --- | --- | --- | --- | --- | --- | --- | --- |
|  | DICE | MHD | ASD | DICE | MHD | ASD | DICE | MHD | ASD | DICE | MHD | ASD |
| Brain_Tech | 0.961 | 8.873 | 0.108 | 0.928 | 5.724 | 0.300 | 0.911 | 7.114 | 0.347 | 0.933 | 7.237 | 0.252 |
| FightAutism | 0.960 | 9.233 | 0.110 | 0.929 | 5.678 | 0.300 | 0.911 | **6.678** | 0.341 | 0.933 | 7.196 | 0.250 |
| OxfordIBME | 0.960 | **8.560** | 0.112 | 0.927 | **5.495** | 0.307 | 0.907 | 6.759 | 0.353 | 0.931 | **6.938** | 0.257 |
| QL111111 | 0.959 | 9.484 | 0.114 | 0.926 | 5.601 | 0.307 | 0.908 | 7.028 | 0.353 | 0.931 | 7.371 | 0.258 |
| Proposed | **0.961** | 9.293 | **0.107** | **0.931** | 5.741 | **0.292** | **0.912** | 7.111 | **0.332** | **0.935** | 7.382 | **0.244** |

3 Note that the best values are highlighted in bold font.

#### 3.3.3. Component analysis of MixACB

We analyzed the effect of the mix ratio on model segmentation performance. Table 5 shows that segmentation accuracy reaches the highest value when the mix ratio is set to 3:1. Then, we further performed an ablation test to verify the effectiveness of each part of the proposed MixACB, as shown in Table 6. The segmentation accuracy was improved with large variations when using different 3D-ACs alone. Moreover, when these 3D-ACs were mixed in scales for a MixACB design, the model was able to achieve the best performance in both DICE and MHD metrics.

**Table 5.** Ablation study performed by comparing the segmentation accuracy in different mix ratios by 2-fold cross validation.

| Mix<br>Ratio | CSF |  | GM |  | WM |  | AVG |  |
| --- | --- | --- | --- | --- | --- | --- | --- | --- |
|  | DICE | MHD | DICE | MHD | DICE | MHD | DICE | MHD |
| 1:0 | 0.952±0<br>.010 | 0.261±0<br>.024 | 0.922±0<br>.008 | 0.604±0<br>.045 | 0.906±0<br>.014 | 0.453±0<br>.041 | 0.927±0<br>.008 | 0.440±0<br>.033 |
| 1:1 | 0.953±0<br>.008 | 0.258±0<br>.023 | 0.921±0<br>.007 | 0.605±0<br>.045 | 0.905±0<br>.013 | 0.455±0<br>.040 | 0.926±0<br>.007 | 0.439±0<br>.033 |
| 3:1<br>(proposed) | **0.953±0**<br>.008 | **0.254±0**<br>.022 | **0.923±0**<br>.007 | **0.601±0**<br>.047 | **0.907±0**<br>.015 | **0.452±0**<br>.043 | **0.928±0**<br>.008 | **0.436±0**<br>.034 |
| 5:1 | 0.953±0<br>.009 | 0.257±0<br>.025 | 0.922±0<br>.008 | 0.601±0<br>.047 | 0.907±0<br>.015 | 0.452±0<br>.042 | 0.926±0<br>.008 | 0.437±0<br>.034 |

Note that the best values are highlighted in bold font.

**Table 6.**
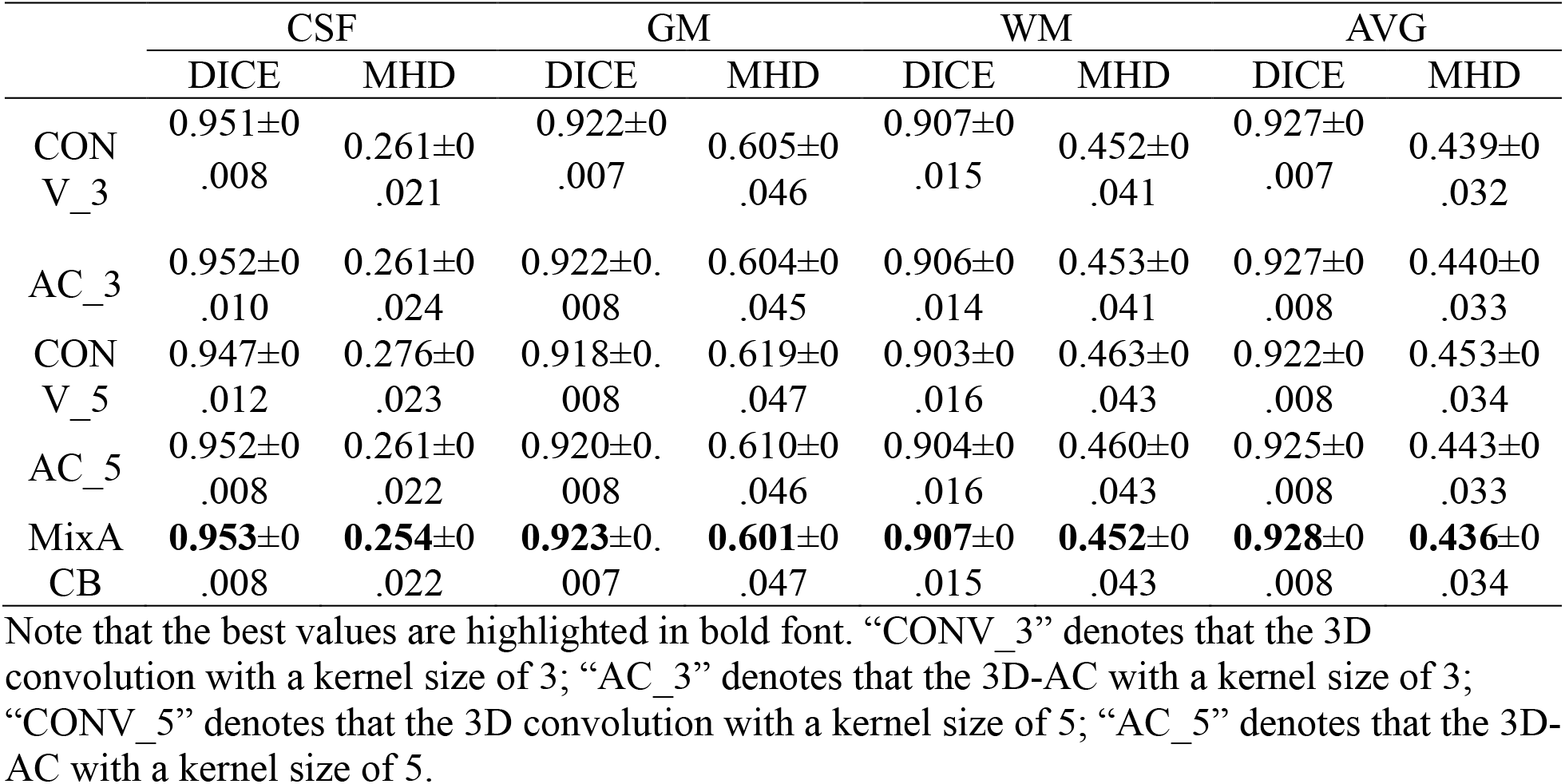
Component analysis of MixACB by 2-fold cross validation.

#### 3.3.4. Impact of overlapping step sizes

We further performed experiments to evaluate the effectiveness of the MixACB on overlapping step sizes, which controls the trade-off between accuracy and inference time. Based on 2-fold cross-validation, which has been done previously, we tested the overlapping impact when the step size is set to 4, 8, 16, and 32 on the DU-Net in the proposed 3D-MASNet framework. Fig. 6A and Fig. 6B present the changes in the segmentation performance in terms of DICE and MHD, respectively, for different overlapping step sizes. Fig. 6C presents the changes in the average number of inference patches for different overlapping step sizes. We found that a step size of 8 is a reasonable choice for achieving fast and accurate results.

**Figure 6.**
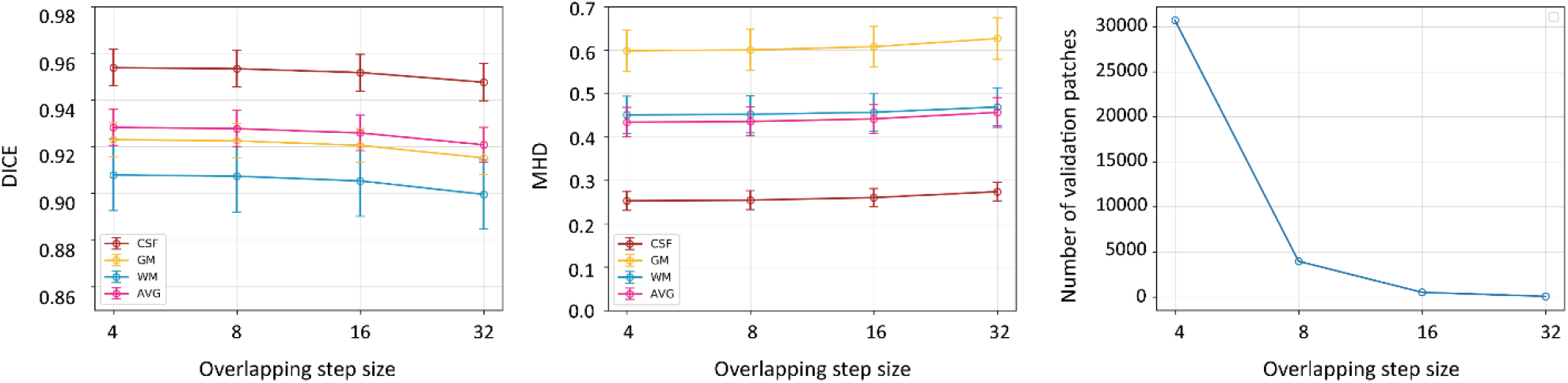
Changes in segmentation performance in terms of DICE (A) and MHD (B) with respect to different overlapping step sizes on 10 subjects during inference, where 2-fold cross-validation is used. (C) Changes of the average number of the 10 subjects’ patches with respect to different overlapping step sizes during inference.

## 4. Discussion

Instead of designing a new network architecture to segment the brain images of 6-month-old infants, we proposed a 3D-MASNet framework by replacing the standard convolutional layer with MixACB on an existing mature network and reduced model parameters and computations by equivalently performing fusion during the inference phase. The experimental results revealed that the MixACB significantly improved the performance of several CNNs by a considerable margin, in which DU-Net with MixACB showed the best average segmentation accuracy. The proposed framework obtained the highest average DICE of 0.935 and lowest ASD of 0.244, which ranked first among all 30 teams on the validation dataset of the iSeg-2019 Grand Challenge. In addition, the CC-3D-FCN model showed the largest improvement, which indicates that a simple model could achieve relatively better performance by implementing our convolution design.

### 4.1. Effectiveness of the MixACB on improving segmentation accuracy

The wide improvement in the segmentation accuracy of different models by the MixACB is derived from several aspects. First, the mixed-scale design of MixACB enables the network to collect multiscale details of local features with different receptive fields, facilitating the integration of coarse-to-fine information inside the input patches at low-to-high semantic levels. Second, the isointense intensity distribution and heterogeneous tissue contrasts hamper effective feature extraction in baby brain images. The mix rate at 3:1 of feature maps for multi-scale kernel size enriches small receptive fields with enough detail features while enabling large receptive fields for capturing coarse global features. We also employed the 3D-AC inside the MixACB by adding multiple orthogonal 3D asymmetric convolutional layers to emphasize informative feature patterns in the central place (Fig. 3). Meanwhile, the asymmetric design is also shown robustness to image rotational distortion (Ding et al., 2019), which may help the network cope with the residual head motion of infants, even though these images have been linearly aligned to standard space. Third, the significant improvable performance of MixACB on various segmentation networks (Table 3) indicates that inter-layer architecture design may not be sufficient for multi-scale information fusion. Notably, besides providing better performance than the previous networks, 3D-MASNet is also more efficient than the baseline models, requiring fewer model parameters once its parameters were fused in the inference phase. For example, the baseline DU-Net’s number of parameters is 2,492,795, while the corresponding 3D-MASNet’s number of parameters is reduced to 2,341,141 during the inference phase.

### 4.2. Well-designed convolution operations

In recent years, researchers have begun to shift their interests from macro network layout to micro neuron units by studying specific convolution operators rather than touching the overall network. Previous works have proposed several advanced convolution operators by combining well-designed filters, such as pyramidal convolution (PyConv), dynamic group convolution (DGC), and asymmetric convolution block (ACB). PyConv employs multiple kernels in a pyramidal way to capture different levels of image details (Duta et al., 2020); DGC equips a feature selector for each group convolution conditioned on the input images to adaptively select input features (Su et al., 2020); ACB introduces asymmetry into 2D convolution to power up the representational power of the skeleton part of the kernel (Ding et al., 2019). These operators implanted into existing mature networks have achieved better performance on image classification or semantic segmentation tasks than in original networks. Due to the “easy-to-plug-in” property, this type of design could be conveniently adopted in various advanced CNNs and avoided high cost of network re-designing. However, these studies mainly concentrate on natural image tasks, few were applied to infant brain segmentation tasks. Here, we design a novel convolution block by combining three basic characteristics including 3D spatial convolution, group convolution containing mixed-scale of kernel sizes, and asymmetry convolution. Due to blurred image appearance, large individual variation of brain morphology, and limited labeled sample sizes, we emphasize that effective and robust feature extraction, especially in a plug-and-play form, is essential for the infant brain segmentation task. Nevertheless, exhausting the combination of various convolution designs is beyond the scope of the article.

### 4.3. Limitations and future directions

The current study has several limitations. First, the patching approach may cause spatial consistency loss near boundaries. Although we adopted a small overlapping step size to relieve this issue, it is necessary to consider further integrating guidance from global information. Second, the small sample sizes of infant-specific datasets limit the generalizability of our method for babies across MRI scanners and acquisition protocols. Further validation on large samples is needed. Third, image indexes, such as the fractional anisotropy derived from diffusion MRI, contain rich white matter information (Liu et al., 2007), which could be beneficial for insufficient tissue contrast (Nie et al., 2019; Zhang et al., 2015). Importantly, determining how to leverage mixed-scale asymmetric convolution to enhance specific model features needs to be further explored. Fourth, we only explored the effectiveness of MixACB when input feature maps are split into 2 groups. Further combination configurations of convolutional kernel sizes and mix ratios are warranted.

## 5. Conclusion

In this paper, we proposed a 3D-MASNet framework for brain MR image segmentation of 6-month-old infants, which ranked first in the iSeg-2019 Grand Challenge. We demonstrated that the designed MixACB could easily migrate to various network architectures and enable performance improvement without extra inference-time computations. This work shows great adaptation potential for further improvement in future studies on brain segmentation.

## Acknowledgments

The study was supported by the National Natural Science Foundation of China (Nos. 31830034, 82021004 and 81801783), Changjiang Scholar Professorship Award (T2015027), and the China Postdoctoral Science Foundation (2020TQ0050 and 2022M710433).

## Data and Code Availability Statement

The 6-months-old infant brain MRI data was publicly offered by the iSeg-2019 (http://iseg2019.web.unc.edu/) organizers.

Codes developed for the proposed segmentation algorithm are released at https://github.com/RicardoZiTseng/3D-MASNet.

